# Multiplex genomic recording of enhancer and signal transduction activity in mammalian cells

**DOI:** 10.1101/2021.11.05.467434

**Authors:** Wei Chen, Junhong Choi, Jenny F. Nathans, Vikram Agarwal, Beth Martin, Eva Nichols, Anh Leith, Choli Lee, Jay Shendure

**Author notes:** These authors contributed equally to this work (Wei Chen, Junhong Choi).

## Abstract

Measurements of gene expression and signal transduction activity are conventionally performed with methods that require either the destruction or live imaging of a biological sample within the timeframe of interest. Here we demonstrate an alternative paradigm, termed ENGRAM (<u>EN</u>hancer-driven <u>G</u>enomic <u>R</u>ecording of transcriptional <u>A</u>ctivity in <u>M</u>ultiplex), in which the activity and dynamics of multiple transcriptional reporters are stably recorded to DNA. ENGRAM is based on the prime editing-mediated insertion of signal- or enhancer-specific barcodes to a genomically encoded recording unit. We show how this strategy can be used to concurrently genomically record the relative activity of at least hundreds of enhancers with high fidelity, sensitivity and reproducibility. Leveraging synthetic enhancers that are responsive to specific signal transduction pathways, we further demonstrate time- and concentration-dependent genomic recording of Wnt, NF-κB, and Tet-On activity. Finally, by coupling ENGRAM to sequential genome editing, we show how serially occurring molecular events can potentially be ordered. Looking forward, we envision that multiplex, ENGRAM-based recording of the strength, duration and order of enhancer and signal transduction activities has broad potential for application in functional genomics, developmental biology and neuroscience.

## Introduction

During development, a modest number of core signaling pathways and gene regulatory modules are leveraged to program a precise spatiotemporal unfolding of programs of cell differentiation, proliferation, morphogenesis and tissue patterning^1^. Across species, differences in how these conserved pathways and modules are used underlies an incredible diversity of organismal form and function. Within species, genetic differences and environmental effects are presumed to influence these core modules in specific developmental or homeostatic contexts, giving rise to both natural phenotypic variation as well as myriad disease states.

How do we capture the activity of these pathways and modules? Measurements of gene expression and signal transduction activity are conventionally performed with methods that require either the destruction or live imaging of a biological sample. These include RNA sequencing (RNA-seq), which measures the global transcriptional state of a system; massively parallel reporter assays (MPRAs), which use sequencing to measure the relative ability of members of a library of DNA fragments to act as enhancers of transcriptional activity in a controlled context^2^; and fluorescent probes and reporters, which track the dynamics of specific signaling pathways in living systems^3^.

These classes of methods are remarkably useful and yet limited in key ways. For example, with RNA-seq, individual samples provide only static snapshots of cell state, such that the temporal dynamics of gene expression must be pieced together by inference with a resolution that is limited by sampling density. Sequencing-based reporter assays are also destructive and static. Although time-series MPRAs can successfully define the temporal dynamics of enhancer activity^4^, such studies are similarly limited by inference and sampling density. Fluorescent probes and reporters are better positioned to capture temporal dynamics, but require that the biological system be physically transparent, at least for live imaging, and are limited in terms of multiplexibility. Overall, there remains a need for a means of capturing signaling and gene regulatory activity that is at once quantitative, reproducible, non-destructive, multiplexable, applicable to physically opaque biological systems and capable of integrating the temporal dynamics of large numbers of signals.

DNA is the natural medium for biological information storage, and is easily “read” through sequencing. To date, a variety of enzymatic systems have been used to alter primary DNA sequence in a signal-responsive manner, most prominently site-specific recombinases (SSRs) and CRISPR-Cas9 genome editing^5^. A classic example is the use of Cre/Lox mice for lineage tracing, wherein the Cre SSR is expressed under the control of a developmentally-specific promoter or enhancer. Cre-mediated recombination at a target locus excises a “stop” sequence, resulting in expression of a fluorescent reporter. The recording event is irreversible, *i*.*e*. the descendants of those cells in which the DNA rearrangement occurred continue to express the reporter regardless of whether the gating regulatory element remains active or not. CRISPR-Cas9 and its derivatives can similarly be used to achieve signal-specific recording. For example, in the proof-of-concept of the CAMERA system^6^, it was shown that expression of a Cas nuclease or base editor could be made dependent on the presence of specific small molecules, light or Wnt signaling. As such, edits mediated by the Cas enzyme served to record those signals.

However, a fundamental limitation of these systems is that they are sharply limited with respect to multiplexibility -- that is, the number of independent signals that can be concurrently recorded. For example, in the case of SSRs or the CAMERA proof-of-concept, enhancers are used to selectively drive the enzyme that mediates an alteration in DNA sequence. However, in this framing, each signal requires its own enzyme, and it is difficult to imagine how more than a handful of independent signals could be concurrently recorded within the same cell or population of cells, let alone how extensive, concurrent recording of large numbers of biological signals could be achieved throughout the development of a multicellular organism.

To address this gap, we developed a new framework for multiplex transcriptional recording, which we term ENGRAM (<u>EN</u>hancer-driven <u>G</u>enomic <u>R</u>ecording of transcriptional <u>A</u>ctivity in <u>M</u>ultiplex). In brief, ENGRAM relies on enzymatic release^7–10^ of prime editing guide RNAs (pegRNAs)^11^ from synthetic transcripts driven by *cis*-regulatory-element (CRE)-coupled Pol-II promoters. Each pegRNA programs the insertion of a specific barcode to a genomically-encoded recording locus (“DNA Tape”). Because each CRE is coupled to a distinct pegRNA-encoded insertion, multiple ENGRAM recorders can operate in parallel, all relying on the same prime editing enzyme and all competing to write to the same DNA Tape.

## Results

### Development and evaluation of ENGRAM 1.0

An ideal DNA-based transcriptional recorder would “log” the production of specific transcripts, *cis-*regulatory activities and/or signal transduction pathways, via specific changes to the primary sequence of a genomic “recorder locus”. Recently developed approaches to such logging of transcripts exploit CRISPR-Cas spacer acquisition^12-17^. However, due to a reliance on host factors, this strategy presently remains limited to prokaryotic systems. In seeking to develop a DNA-based recorder for mammalian systems, we were inspired by reporter assays, an established approach wherein a *cis-*regulatory element (CRE) of interest is positioned upstream of a minimal promoter (minP) and reporter gene (*e*.*g*. luciferase). Reporter assays are amenable to extensive multiplexing, as the reporter can include a transcribed barcode that is linked to the CRE, resulting in the MPRA^18^. However, as noted above, MPRAs depend on targeted RNA-seq of the barcodes, which is destructive and static.

Nonetheless, we reasoned that the basic MPRA architecture, *i*.*e*. a library of synthetic or natural enhancers positioned upstream of a minimal promoter, might be coupled to the expression of a library of “writing units”, in the form of guide RNAs (**Figure 1a**). Each CRE would be associated with a specific guide RNA, whose expression would trigger a specific edit. A first challenge to this scheme is that on their own, CRISPR-Cas9 single guide RNAs (sgRNAs) program the location of editing but not the edit itself. As such, each CRE of interest would require its own target, rather than recording to a shared DNA Tape. However, this is potentially addressable with prime editing^11^, as a pegRNA programs both the site of editing and the edit itself. Specifically in ENGRAM, each CRE is linked to a pegRNA encoding a specific insertion to a common DNA Tape.

**Figure 1.**
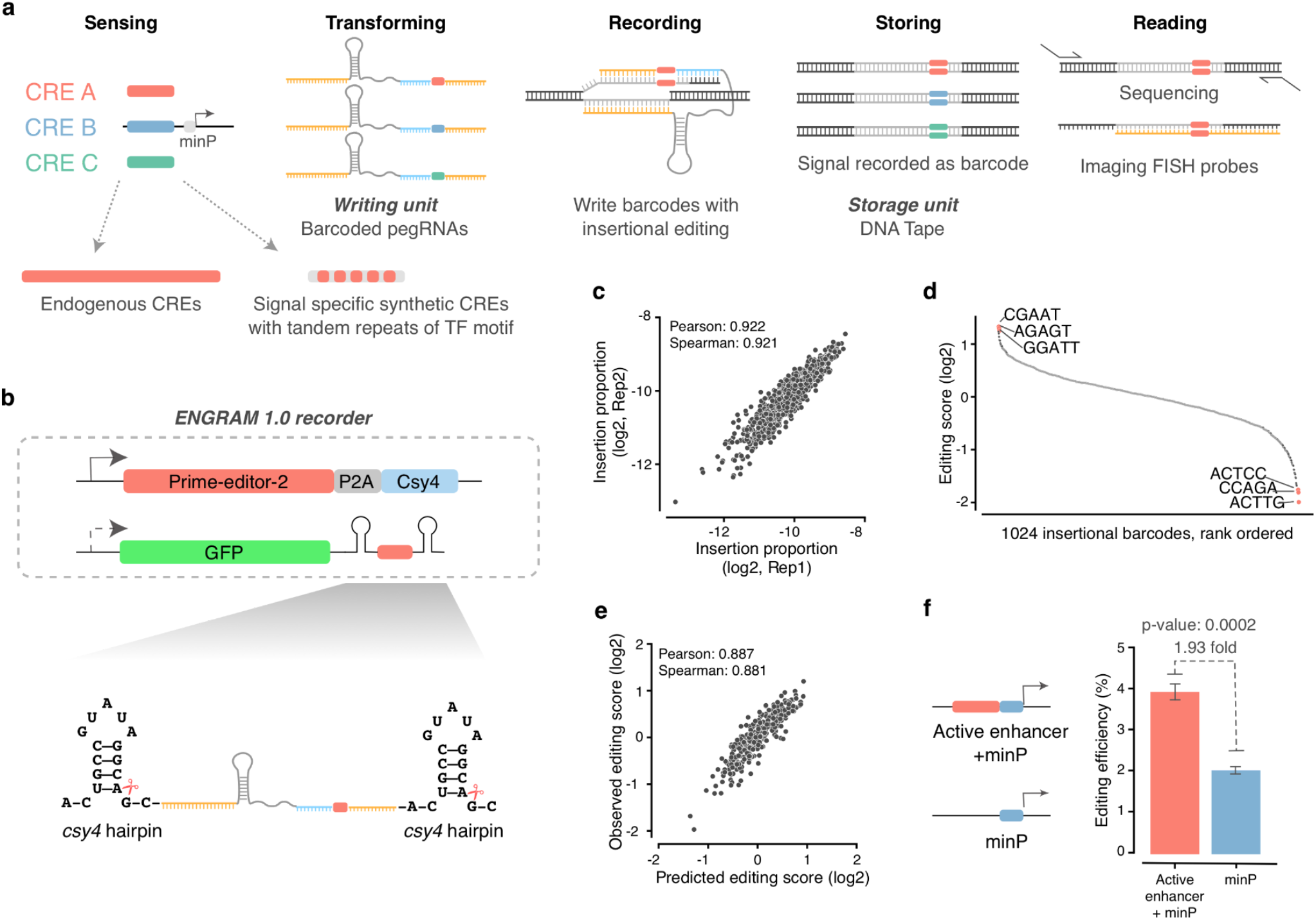
ENhancer-driven Genomic Recording of transcriptional Activity in Multiplex (ENGRAM). **(a)** Schematic of ENGRAM. Endogenous or synthetic *cis-*regulatory elements (CREs) drive activity-dependent transcription of a prime editing guide RNA (pegRNA) encoding a CRE-specific insertion. The insertion is written to a natural or engineered recording site within genomic DNA (“DNA Tape”). The recorded signal can subsequently be recovered by DNA sequencing or, potentially, by fluorescent *in situ* hybridization (FISH). **(b)** Schematic of the ENGRAM 1.0 recorder. A pegRNA writing unit is flanked by *csy4* hairpins and embedded within the 3’ UTR of a Pol-2-driven GFP mRNA. PE2 and Csy4 are constitutively expressed from a separate locus. Csy4 cleaves at the *csy4* hairpins and releases the active pegRNA. **(c)** 1024 5N barcodes inserted to *HEK3* locus by pegRNAs (ENGRAM 1.0 design) driven by a constitutive Pol-2 PGK promoter. Log-scaled insertion proportions (calculated as the proportion of edited *HEK3* sites with a given insertion) are highly correlated between transfection replicates. **(d)** Range of editing scores (ES) for 5N insertions. ES are calculated as (genomic reads with specific insertion/total edited *HEK3* reads)/(plasmid reads with specific insertion/total plasmid reads), plotted here in rank order on a log2-scale. **(e)** A linear lasso regression model trained on these data with one-hot encoded single and dinucleotide content of the 5-mer predicts insertional efficiencies with reasonably high accuracy. Samples were split with 724 barcodes in a training set and 300 barcodes in a test set. The model was trained with 10-fold cross validation on the training set, and then used to predict the test set. **(f)** A schematic of the constructs used for the two pools of ENGRAM 1.0 recorders is shown on the left, and the observed editing efficiency for each pool on the right. Briefly, a pool of 13 enhancers known to be active in this cell line, cloned upstream of minP and driving a pool of pegRNAs encoding insertion of a 5N degenerate sequence to *HEK3*, was 1.93-fold more active than a control construct bearing minP alone. Error bars correspond to standard deviations across 3 transfection replicates. P-value = 0.0002 (t-test).

A second challenge is that in order to be appropriately processed, transcripts for most translated genes, including CRE-minP-driven reporter transcripts, are made by RNA polymerase II (Pol-2), whereas small untranslated RNAs, including guide RNAs, are made by RNA polymerase III (Pol-3). To address this, we leveraged the CRISPR endoribonuclease Csy4 (also known as Cas6f), which recognizes and cuts at the 3’ end of 17-bp RNA hairpins (*csy4*)^7–10^. Constitutive expression of Csy4, together with CRE-activity-dependent expression of *csy4*-pegRNA-*csy4*, should result in a liberated pegRNA (**Figure 1b**). To further facilitate transcription and stabilize the mRNA^19^, we embedded *csy4*-pegRNA-*csy4* within the 3’ untranslated region (UTR) of a GFP transcript (**Figure 1b**). Thus, the ENGRAM 1.0 recorder architecture consists of a CRE-coupled Pol-2 promoter followed by a GFP transcript with *csy4*-pegRNA-*csy4* embedded in its 3’ UTR.

To benchmark the activity of pegRNAs released from Pol-2 transcripts, we compared an ENGRAM 1.0 recorder driven by a constitutive Pol-2 promoter (PGK) to a conventional, U6-driven pegRNA. In both cases, the pegRNAs target the endogenous HEK293 target 3 (*HEK3*) locus and are designed to insert three nucleotides (CTT)^11^. These constructs were separately transiently transfected to monoclonal HEK293T cells constitutively expressing Prime-Editor-2 (PE2) and Csy4. Five days after transfection, we harvested genomic DNA, and then PCR-amplified and sequenced the *HEK3* locus. We observed comparable, reproducible efficiencies of CTT insertion between the ENGRAM 1.0 and U6 recorders (mean 5.9% and 5.3% across three replicates, respectively; **Supplementary Figure 1a**).

To evaluate whether prime editing-mediated insertions could be used as barcodes, we next sought edit the endogenous *HEK3* target via transient transfection with a pool of plasmids encoding PGK-driven ENGRAM 1.0 pegRNAs bearing a degenerate 5N region in place of CTT. Although editing efficiencies were lower than for the CTT insertion on average, all 1,024 potential 5-bp insertions were observed at the *HEK3* locus at appreciable, reproducible frequencies (**Figure 1c**; **Supplementary Figure 1a-d**). After normalizing for their abundance in the plasmid pool, the insertional efficiencies of individual 5-mers were generally balanced, with 93% falling within a 4-fold range (**Figure 1d**). To assess the sequence determinants of differences in insertional efficiency, we performed linear lasso regression with one-hot single and dinucleotide encoding of the 5-mer; the resulting model was reasonably accurate in predicting these biases (**Figure 1e**; **Supplementary Figure 1e**).

Finally, we sought to apply ENGRAM 1.0 as an enhancer-driven recorder, by replacing the constitutive PGK promoter with a CRE-minP architecture. We selected thirteen 170-bp sequences previously shown to have enhancer activity in K562 cells^18^. We cloned these upstream of minP and ENGRAM 1.0 pegRNAs encoding a degenerate 5N insertion. We then compared the editing efficiency of the pool of enhancer-driven recorders vs. a pool of negative controls (minP with no upstream enhancer) via their transient transfection to K562 cells constitutively expressing both PE2 and Csy4. We successfully recorded enhancer-activated barcode insertions with a collective efficiency of 3.9%, 1.93-fold higher than the editing efficiency of pegRNAs driven by minP alone (**Figure 1f**). Overall, these results suggest that ENGRAM-based recording can work. However, the signal-to-noise ratio was considerably more modest than we had hoped for. We speculated that this was due in part to the accumulation of background edits due to constitutive expression of Csy4, which may also have reduced growth of the monoclonal cell line due to cytotoxic effects^10^.

**Supplementary Figure 1.**
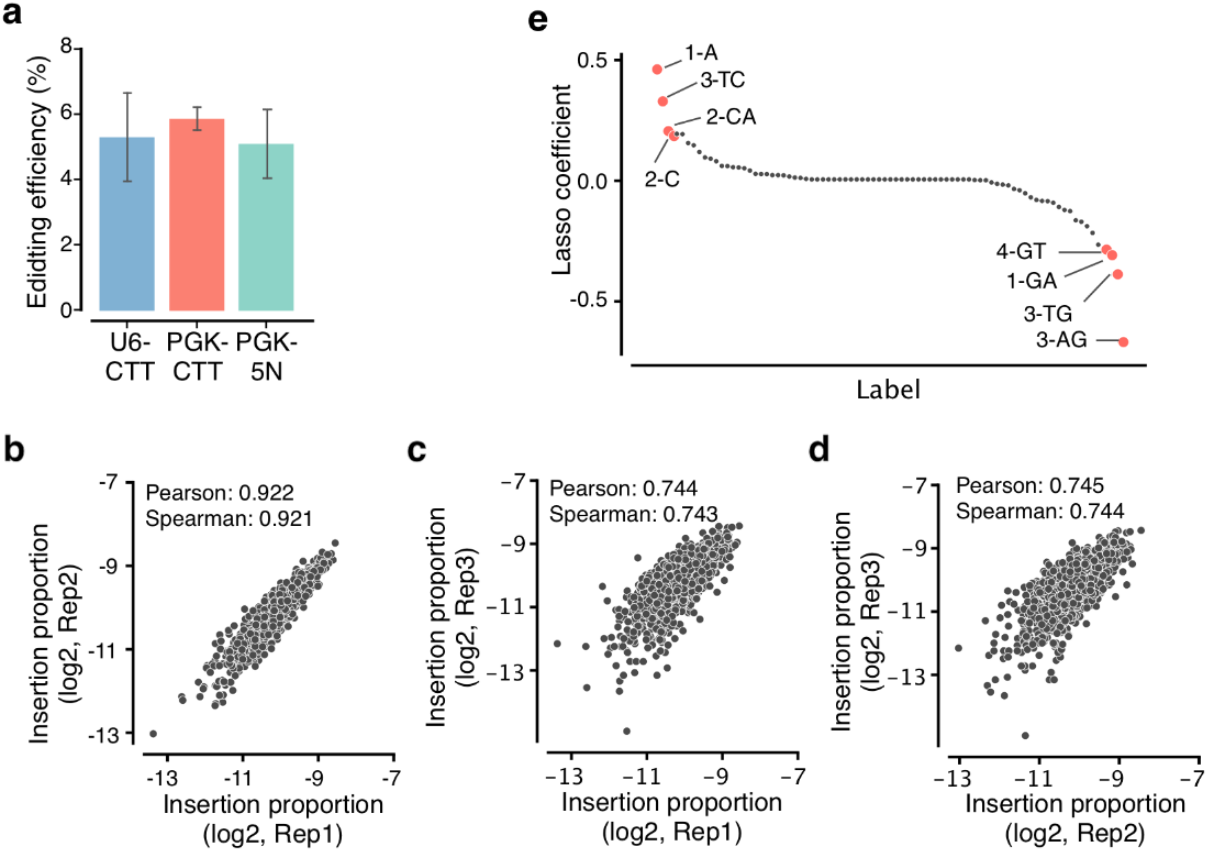
The ENGRAM 1.0 recorder installs barcodes with reasonable efficiency and reproducibility. **(a)** Across three transfection replicates, the ENGRAM 1.0 recorder driven by a constitutive Pol-2 PGK promoter (PGK-CTT) exhibited comparable efficiency for inserting CTT at the *HEK3* locus to a U6-driven CTT-pegRNA (U6-CTT). A similar ENGRAM 1.0 recorder driving a degenerate 5N insertion rather than CTT (PGK-5N) exhibited comparable efficiency as well. In the K562 cell line in which this experiment was performed, PE2 and Csy4 were constitutively expressed. **(b-d)** Reproducibility of the relative proportions of 1024 5N barcodes installed by ENGRAM 1.0 driven by the constitutive Pol-2 PGK promoter. Log-scaled insertion proportions (calculated as the proportion of edited *HEK3* sites with a given insertion) were well correlated between pairs of transfection replicates. **(e)** A linear lasso regression model trained on these data with one-hot encoded single and dinucleotide content predicts insertional efficiencies with reasonably high accuracy. Samples were split with 724 barcodes in a training set and 300 barcodes in a test set. The model was trained with 10-fold cross validation on the training set, and then used to predict the test set. Plotted here are the rank-ordered coefficients of the linear lasso regression. Positional information of single nucleotides and dinucleotides were used as input features for training. The top 4 and bottom 4 coefficients are annotated (*e*.*g*. 1-A and 3-TC mean A at first nucleotide or TC dinucleotide starting at position 3, respectively).

### Multiplex recording of enhancer activity with ENGRAM 2.0

To reduce the background accumulation of edits to the DNA Tape, we designed a new ENGRAM architecture in which the GFP ORF is replaced by Csy4 ORF, and Csy4 is no longer constitutively expressed (**Figure 2a**). In this recorder design, termed ENGRAM 2.0, the expression of Csy4 and the pegRNA are <u>both</u> dependent on enhancer activity. To evaluate whether these modifications reduce background recording, we tested ENGRAM 1.0 vs. 2.0 in the absence of any enhancer, *i*.*e*. minP alone driving pegRNAs that programmed a degenerate 5N insertion to the endogenous *HEK3* target locus. Transiently co-transfecting these constructs into HEK293T cells with either PE2-Csy4 plasmid (for ENGRAM 1.0) or PE2 plasmid (for ENGRAM 2.0) in triplicate, we indeed observed a 2.8-fold reduction in background recording with ENGRAM 2.0 relative to ENGRAM 1.0 (mean 1.4% for ENGRAM 1.0 → 0.5% for ENGRAM 2.0, 3 days post-transfection) (**Figure 2b**).

**Figure 2.**
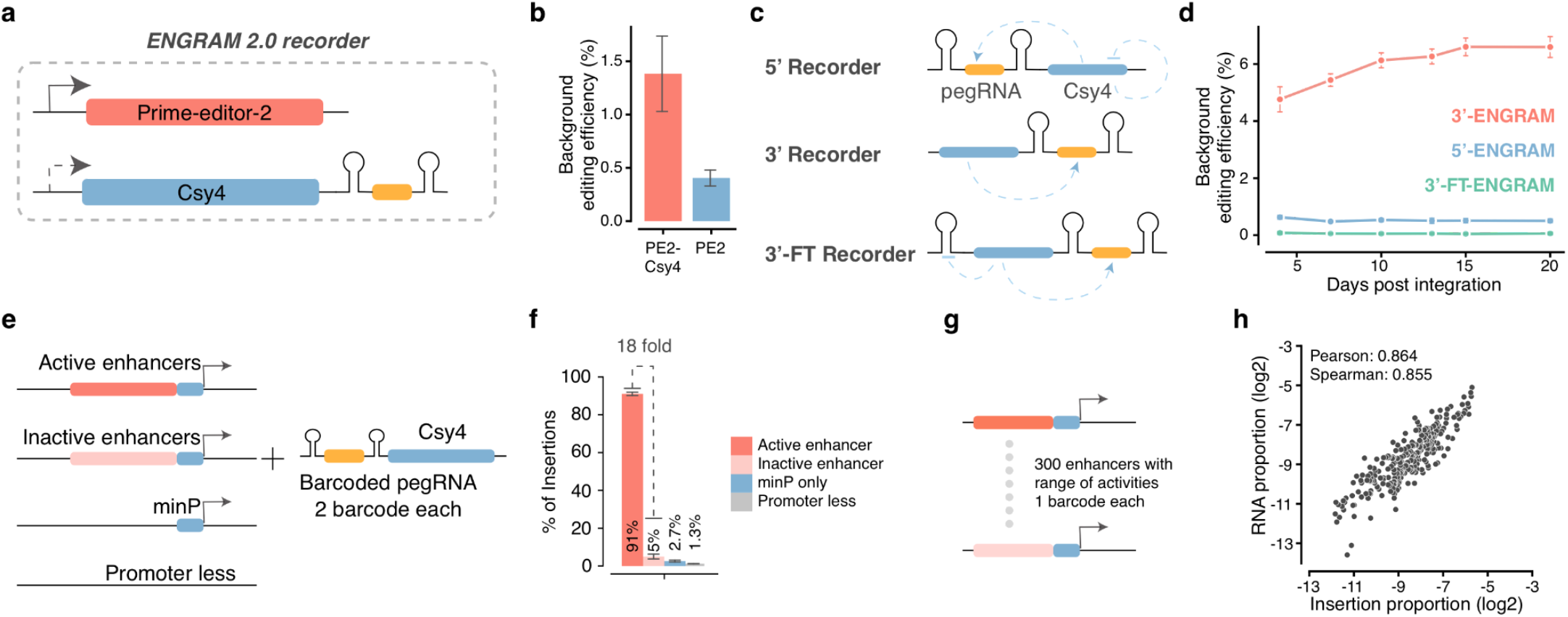
Development and benchmarking of ENGRAM 2.0 recorders. **(a)** In ENGRAM 2.0, Csy4 is encoded by the same transcript that encodes the *csy4* hairpin*-*flanked pegRNA. This is expected to reduce background activity, relative to ENGRAM 1.0, where Csy4 was constitutively expressed. **(b)** ENGRAM 2.0 exhibits lower levels of background recording than ENGRAM 1.0. Measurements are for minP alone driving pegRNAs programming a degenerate 5N insertion to the *HEK3* locus in triplicate, 3 days post-transfection. Error bars correspond to standard deviations across 3 transfection replicates. **(c)** Three versions of ENGRAM 2.0 with the *csy4* hairpin-flanked pegRNA embedded in the 5’ or 3’ UTR of a transcript encoding Csy4, or 3’ ENGRAM 2.0 with an additional *csy4* hairpin in the 5’ UTR in order to impose auto-regulatory negative feedback on Csy4 levels. **(d)** All three ENGRAM 2.0 recorders were integrated via piggyBAC into PE2-expressing cells in triplicate, each driving 1,024 5N barcodes with minP. The background editing efficiency was periodically checked over 20 days. Error bars correspond to standard deviations across 3 transfection replicates. **(e)** Benchmarking of ENGRAM 2.0 with enhancers with known activities in a reporter assay. 5’-ENGRAM recorders with active and inactive enhancers upstream of a minP, together with minP-only and promoter-less constructs, were cloned, each driving expression of distinct pegRNA-encoded barcodes. **(f)** Barcodes corresponding to the active enhancer comprised the vast majority (91.1%) of recorded events. Error bars correspond to standard deviations from 3 transfection replicates. **(g-h)** Further benchmarking of ENGRAM 2.0 with 300 enhancers known to have a range of activities in a reporter assay. **(g)** This library was designed such that each enhancer drove expression of a distinct pegRNA-encoded 6-mer insertional barcode. **(h)** alues correspond to the proportion of each barcode read out from the *HEK3* genomic locus (ENGRAM) or from the pegRNAs (MPRA), out of the total. Proportions were corrected for sequence-based insertional biases (details in **Supplementary Figure 2f**). The log-scaled proportions of ENGRAM events recorded to DNA were highly correlated with log-scaled proportions of barcodes measured directly from RNA.

Towards further reducing background, we designed two additional recorders: 5’ ENGRAM 2.0, in which the *csy4* hairpin-flanked pegRNA is embedded within the 5’ (rather than 3’) UTR of the *Csy4* transcript; and 3’-FT ENGRAM 2.0, which contains an additional *csy4* hairpin in its 5’ UTR to create auto-regulatory negative feedback loop on Csy4 levels (**Figure 2c**). Integrating these recorders into HEK293T cells expressing PE2 (PE2(+) HEK293T) cells via piggyBAC and then evaluating background activity as above, the 5’ ENGRAM 2.0 and 3’-FT ENGRAM 2.0 recorders respectively exhibited 12-fold and >100-fold reductions in background activity, relative to 3’ ENGRAM 2.0 (10 days post-transfection; **Figure 2d**; **Supplementary Figure 2a**). Of note, for all three of these integrated ENGRAM 2.0 recorders, the level of background recording plateaued after several days (**Figure 2d**). This suggested to us that the accumulation of background recording events mostly occurs shortly after transfection, potentially due to ORI-driven, plasmid-mediated transcription^18,20^, rather than minP-driven transcription from integrated recorders. However, some degree of accumulation persisted with the 3’ ENGRAM 2.0 recorder, suggesting an additional component of genomically driven background activity.

Although the 3’-FT design exhibited the lowest background activity, we moved forward with the 5’ ENGRAM 2.0 design because its organization facilitates straightforward pairing of CREs and pegRNA-mediated insertions during cloning. To further benchmark it, we cloned a pair of 170-bp sequences previously shown to have either high vs. minimal enhancer activity in K562 cells^18^ upstream of minP, together with minP-only and promoter-less constructs (**Figure 2e**). Each of these four constructs drove pegRNAs designed to insert two distinct 5-bp insertions to the endogenous *HEK3* target locus. An equimolar mixture of these 8 recorder plasmids was introduced via piggyBAC integration into PE2-expressing K562 cells in triplicate. At five days post-transfection, 3.14% of *HEK3* target sites were edited, but over 90% of inserted barcodes were associated with the active enhancer (**Figure 2f**; **Supplementary Figure 2b**). Of note, the 18-fold difference in recorded insertional frequency between the active and inactive enhancer roughly matched the 15-fold difference between them measured by MPRA^18^.

To more generally evaluate whether the enhancer activities recorded by ENGRAM are quantitatively comparable to corresponding measurements made by MPRA, we cloned 300 enhancer fragments^18^ to the 5’ ENGRAM 2.0 construct, each driving a pegRNA encoding the insertion of a unique 6-mer to the *HEK3* locus (**Figure 2g**). Four days after introducing these recorders via piggyBAC to PE2-expressing K56 in triplicate, we separately recovered, amplified and sequenced the *HEK3* locus (from DNA) or the transcribed barcode itself (from RNA). From DNA, we observed an overall editing efficiency of 1.97%, and recovered 298 of 300 barcodes. Both RNA and DNA-based measurements were highly consistent between transfection replicates (**Supplementary Figure 2c-d**). Furthermore, we observed a strong correlation between the recorded activities (ENGRAM; DNA) and the directly measured activities (MPRA; RNA), indicating that the relative transcriptional activities of enhancer reporters can be quantitatively recorded to genomic DNA (**Supplementary Figure 2e**). Correction for sequence-based insertional biases further improved the correlation between enhancer-driven reporters (MPRA) and recorders (ENGRAM) (**Figure 2h**; **Supplementary Figure 2f**).

**Supplementary Figure 2.**
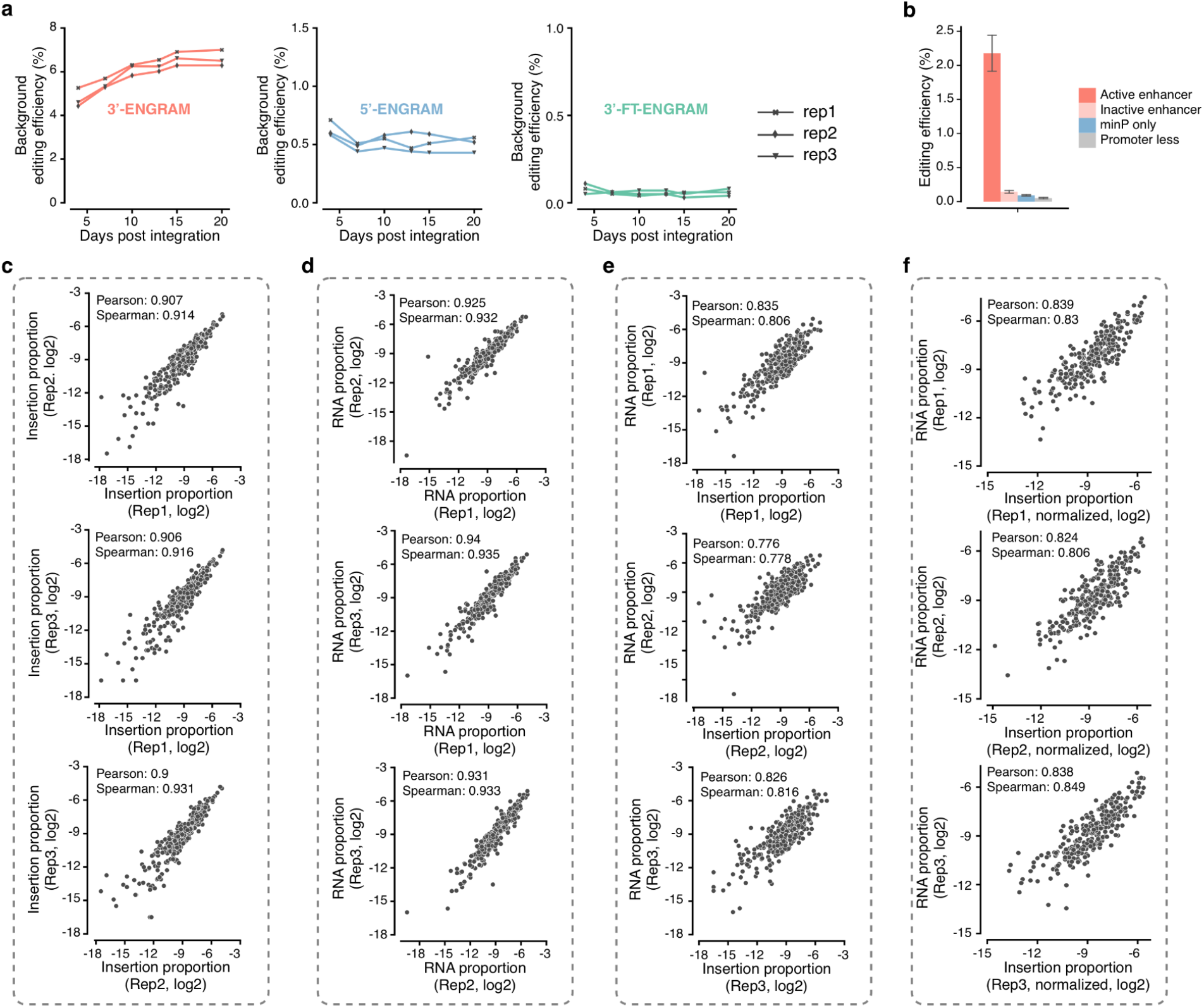
Benchmarking of ENGRAM 2.0 recorders. **(a)** All three ENGRAM 2.0 recorders were integrated via piggyBAC into PE2-expressing cells in triplicate, each driving 1,024 5N barcodes with minP. The background editing efficiency was periodically checked over 20 days. Similar to **Fig. 2b** but with each transfection replicate of each recorder design plotted separately. **(b)** Benchmarking of ENGRAM 2.0 with enhancers with known activities in a reporter assay. 5’-ENGRAM recorders with active and inactive enhancers upstream of a minP, together with minP-only and promoter-less constructs, were cloned, each driving expression pegRNA-encoded barcodes. Barcodes corresponding to the active enhancer were introduced at a much higher rate. Error bars represent standard deviations from 3 transfection replicates. Similar to **Fig. 2f** but showing the editing rate rather than the proportion of barcodes. **(c)** Log-transformed insertion proportions for 300 6-mer barcodes were highly reproducible across transfection replicates. Each value corresponds to the proportion of barcodes read out at the DNA level from the *HEK3* locus. **(d)** Log-transformed RNA proportions for 300 6-mer barcodes were highly reproducible across transfection replicates. Each value corresponds to the proportion of barcodes read out at the RNA level from transcribed pegRNAs. **(e)** The log-scaled proportions of ENGRAM events recorded to DNA were highly correlated with log-scaled proportions of barcodes measured directly from RNA. **(f)** Same as panel **e**, but after correcting for sequence-based insertional biases. In more detail, to determine sequence-based insertional biases for 6-mers, we cloned a library of pegRNAs encoding a degenerate 6N insertion, driven by the highly active PGK promoter. This library was transiently transfected into K562 cells expressing PE2. Genomic DNA was harvested at four days post transfection. For 266 barcodes for which we recovered reads, editing scores (ES) were calculated as (genomic reads with specific insertion/total edited *HEK3* reads)/(plasmid reads with specific insertion/total plasmid reads). To correct the counts of ENGRAM-recorded events for sequence-based insertional bias, we simply divided them by the ES of the corresponding 6-mer insertion. This normalization improved the correlation with reporter-based RNA proportions (panel **e** vs. panel **f**).

### Quantitative recording of signaling pathway activation or small molecule exposure with ENGRAM

We next sought to ask whether ENGRAM could be used to record the intensity or duration of signaling pathway activation or small molecule exposure. For this, we selected several signal-responsive regulatory elements: the Tet Response Element (TRE; activated by doxycycline)^21^, a NF-κB responsive element (activated by TNFα)^22^, and a TCF-LEF responsive element (Wnt signaling pathway; activated by CHIR99021)^23^, each previously used to drive fluorescent reporters in a signal-responsive manner (**Supplementary Table 1**). These signal-responsive sequences were cloned upstream of minP within 5’ ENGRAM 2.0 recorders, with each driving expression of a pegRNA encoding one or two specific insertions to the *HEK3* locus (**Figure 3a**). The three recorders were separately integrated into the genomes of PE2(+) HEK293T cells via piggyBAC in triplicate (for the doxycycline recorder, constitutively expressed reverse tetracycline-controlled transactivator (rtTA) was integrated separately). A 2-fold dilution series of doxycycline, TNFα or CHIR99021 was added to the media of the cell lines into which the relevant recorder had been integrated, and genomic DNA was harvested 48 hours after the onset of exposure.

**Figure 3.**
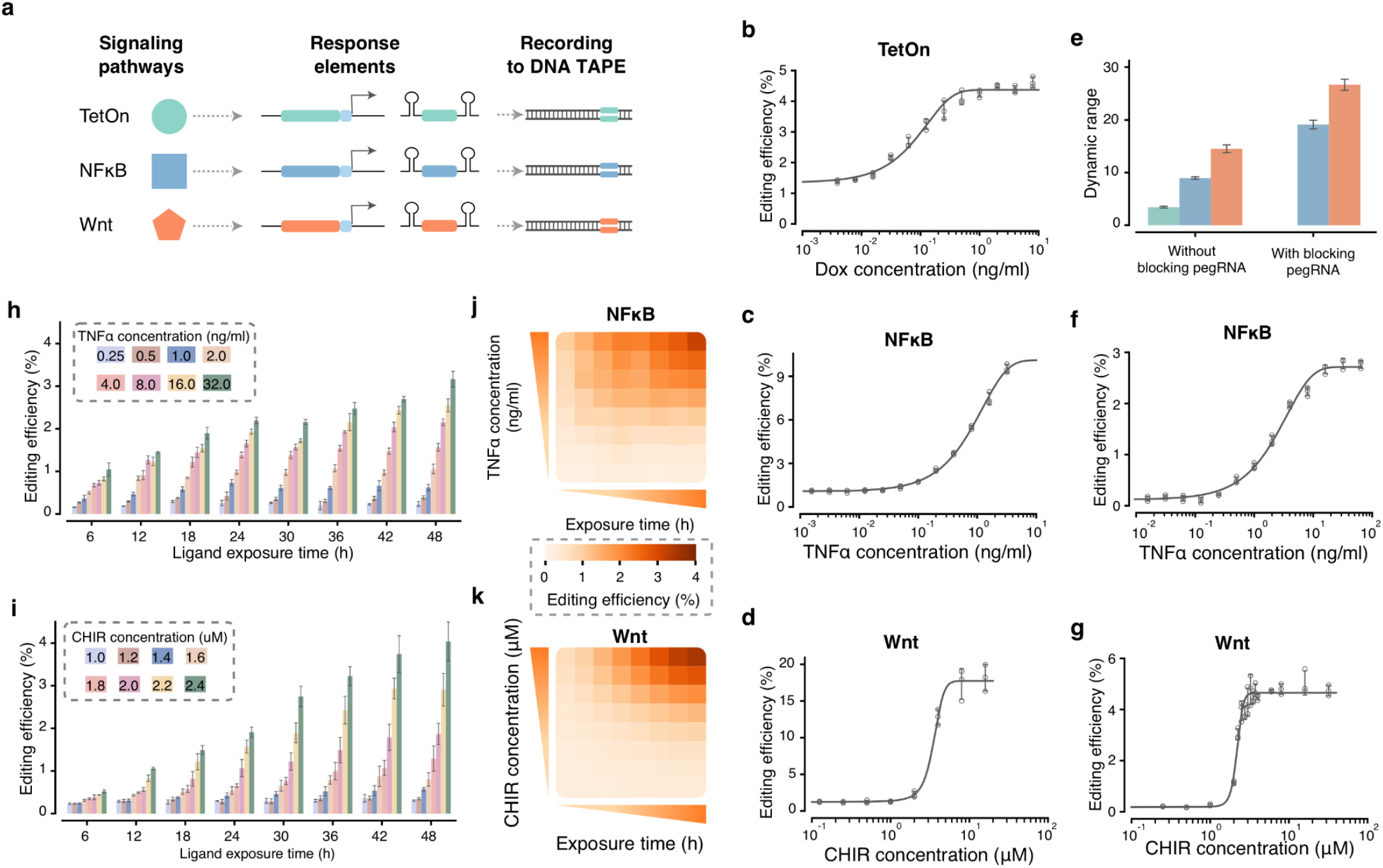
Recording the intensity and duration of signaling pathway activation or small molecule exposure. **(a)** Signal-responsive regulatory elements were used to construct ENGRAM 2.0 recorders for activation by doxycycline (TetON; Tet Response Element), TNFα (a NF-κB responsive element) and CHIR99021 (a TCF-LEF responsive element, responsive to Wnt signaling). **(b-d)** Upon 48 hours of stimulation with the corresponding stimulant, the TetON **(b)**, NF-κB **(c)**, and Wnt **(d)** recorders exhibited dose-dependent levels of recording. These experiments were conducted on separate, polyclonal cell lines, each of which had one recorder integrated via piggyBAC. Cells were exposed to a serial two-fold dilution series of doxycycline **(b)**, TNFα **(c)** or CHIR99021 **(d)**, with starting concentrations of 8 ng/ml, 3.2 ng/ml and 16 µM, respectively. **(e)** Dynamic range observed in signal recording experiments without (left) or with (right) co-transfection of plasmid-encoded, non-targeting, non-integrating blocking pegRNA to suppress background. Colors as in panel **a. (f)** Similar to NF-κB experiment shown in panel **c**, except with co-transfection of plasmid-encoded, non-targeting, non-integrating blocking pegRNA to suppress background. Furthermore, the starting TNFα concentration for the two-fold dilution series was adjusted to 64 ng/ml. **(g)** Similar to Wnt experiment shown in panel **d**, except with co-transfection of plasmid-encoded, non-targeting, non-integrating blocking pegRNAs to suppress background. Additionally, more CHIR99021 concentrations were sampled between 1 to 4 µM. Error bars correspond to standard deviations from 3 stimulus replicates. **(h**,**i)** Histograms showing editing efficiencies resulting from matrix experiment on the NF-κB **(h)** and Wnt **(i)** recorders, in which both stimulant concentrations and durations of exposure were varied (2 recorders x 8 concentrations x 8 durations x 3 replicates = 384 conditions). Error bars correspond to standard deviations from 3 stimulus replicates. **(j**,**k)** Heatmap representation of the same data as in panels **h** and **i**, illustrating the joint dependence of recording levels on the dose and duration of stimulation.

For all three signal-responsive ENGRAM recorders, editing rates at the *HEK3* locus exhibited a strikingly sigmoidal dependence on the log-transformed concentration of the corresponding stimulant (**Figure 3b-d**). This was particularly the case for the Wnt signaling, wherein the corresponding recorder exhibited nearly switch-like behavior across an approximately four-fold range of CHIR99021 concentration (**Figure 3d**). As with previous experiments, each ENGRAM recorder exhibited some degree of non-accumulating, basal recording even in the absence of signal exposure (1.4%; **Supplementary Figure 3a**), potentially due to ORI-driven, plasmid-mediated transcription^18,20^ shortly after transfection, as discussed above. Despite this background, we observed a dynamic range in editing efficiency between background vs. maximal stimulation of 3.3-fold, 9.0-fold and 14.9-fold for the Tet, NF-κB and Wnt recorders, respectively (**Figure 3e**).

In an attempt to further improve signal-to-noise, we remade the NFΔB and Wnt recorder cell lines, but with co-transfection of a non-integrating plasmid expressing a non-targeting pegRNA driven by a U6 promoter. The intent of this non-targeting pegRNA was to compete for PE2 binding and thereby minimize any “pre-integration” activity from pegRNAs encoded by the ENGRAM recorders. Indeed, the plasmid expressing the blocking pegRNA reduced background recording to from ∼1.4% to 0.14% (for NFκB recorder) and 0.22% (for Wnt recorder) (**Supplementary Figure 3b**).

After waiting five days to allow for the plasmid expressing the blocking pegRNA to be diluted out and the ENGRAM recorders to be integrated, we repeated the dilution series experiment, further using this opportunity to adjust the concentrations sampled (*e*.*g*. for Wnt, more densely sampling CHIR99021 concentrations around the inflection point of the dose-response curve; **Figure 3d**,**g**). Although the overall levels of recording were reduced, possibly due to a lower multiplicity of infection of the piggyBAC-based ENGRAM recorder plasmids, the signal-to-noise approximately doubled, with a dynamic range in editing efficiency between background vs. maximal stimulation of 19.0-fold and 27.3-fold for the NF-κB and Wnt recorders, respectively (**Figure 3e-g**).

To explore the dependence of ENGRAM on not only the intensity of signals but also their duration, we performed a matrix experiment on the NF-ΔB and Wnt recorders, varying stimulant concentration as previously but also varying the duration of exposure from 6 to 48 hours (2 recorders x 8 concentrations x 8 durations x 3 replicates = 384 conditions; **Figure 3h-k**). In this experiment, each batch of cells was harvested 24 hours after the removal of stimulants from the media. In the resulting levels of editing, the dependency of the NF-ΔB and Wnt recorders on both the intensity and duration of stimulation was immediately evident (**Figure 3j-k**). For both recorders, even 6 hours of stimulation was sufficient to observe signal in excess of background. However, the NF-ΔB recorder appeared to exhibit faster kinetics than the Wnt recorder (**Figure 3h-i**).

**Supplementary Figure 3.**
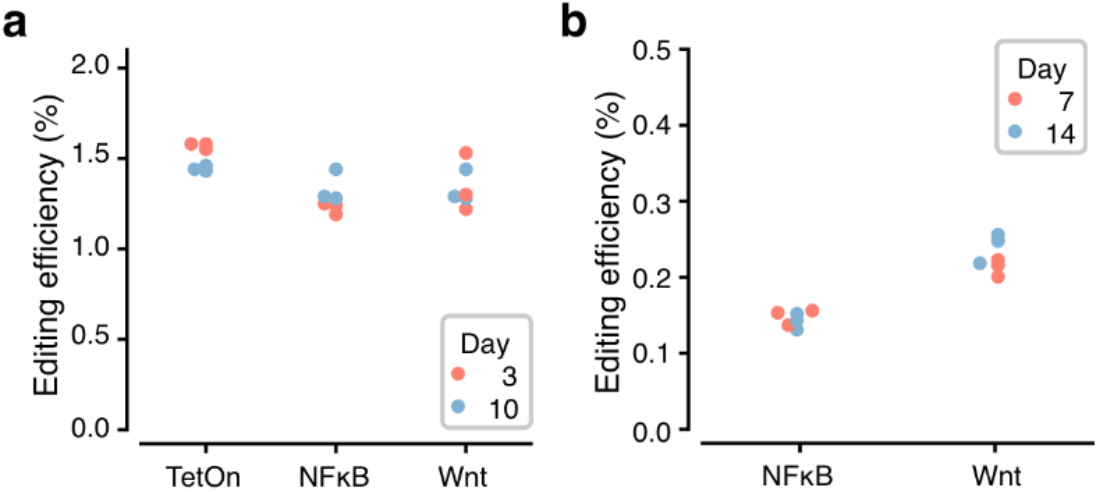
Successful suppression of pre-integration background accumulation. **(a)** As with the enhancer-responsive ENGRAM recorders (**Figure 2d**; **Supplementary Figure 2a**), we observed modest levels of background recording even in the absence of stimulus with the signal-responsive ENGRAM recorders. Once again, this background did not accumulate over time, consistent with the hypothesis that it primarily accumulates shortly after transfection, potentially due to ORI-driven, plasmid-mediated transcription. **(b)** We repeated the construction of the NFκB and Wnt-responsive ENGRAM recorders but with co-transfection of a plasmid-encoded, non-targeting, non-integrating blocking pegRNA expressed from a U6 promoter. This successfully reduced background, which was once again non-accumulating over time, from ∼1.4% to 0.14% (for the NFκB recorder) and 0.22% (for the Wnt recorder).

### Multiplex recording of signaling pathway activity with ENGRAM

We next sought to introduce multiple ENGRAM recorders for different signaling pathways into a single population of cells, to evaluate whether they could be used together, *i*.*e*. competing to write to a shared DNA Tape (**Figure 4a**). In brief, constructs corresponding to the TetON, NF-κB and Wnt recorders were mixed at an equimolar ratio and co-integrated to PE2(+) HEK293T cells. Each recorder drives pegRNA(s) encoding the insertion of one or two distinct, signal-specific barcodes (**Supplementary Table 1**). These cells were exposed to a high concentration of all possible combinations of 0 to 3 stimuli, in triplicate (8 on/off stimulus combinations x 3 replicates = 24 conditions). Harvesting cells after 48 hours of stimulation, we performed PCR amplification and sequencing of the shared DNA tape. As predicted, the abundances of signal-specific barcodes were highly dependent on the precise combination of stimuli applied (**Figure 4b**). Put another way, we observed minimal cross-talk, consistent with the orthogonality of these signaling pathways to one another (**Supplementary Figure 4a**). To push this system further, we performed a separate experiment in which populations of cells bearing all three recorders were exposed to all possible combinations of low, medium or high concentrations of each stimulus (3 concentrations ^ 3 stimuli x 3 replicates = 81 conditions). Once again harvesting cells after 48 hours and reading the DNA Tape, we observe that signal-specific barcodes are introduced at rates correlated with the concentration of the corresponding stimulus (**Figure 4c**; **Supplementary Figure 4b**), further supporting the conclusion that these recorders are able to capture quantitative information on separate channels despite writing to a shared DNA Tape.

**Figure 4.**
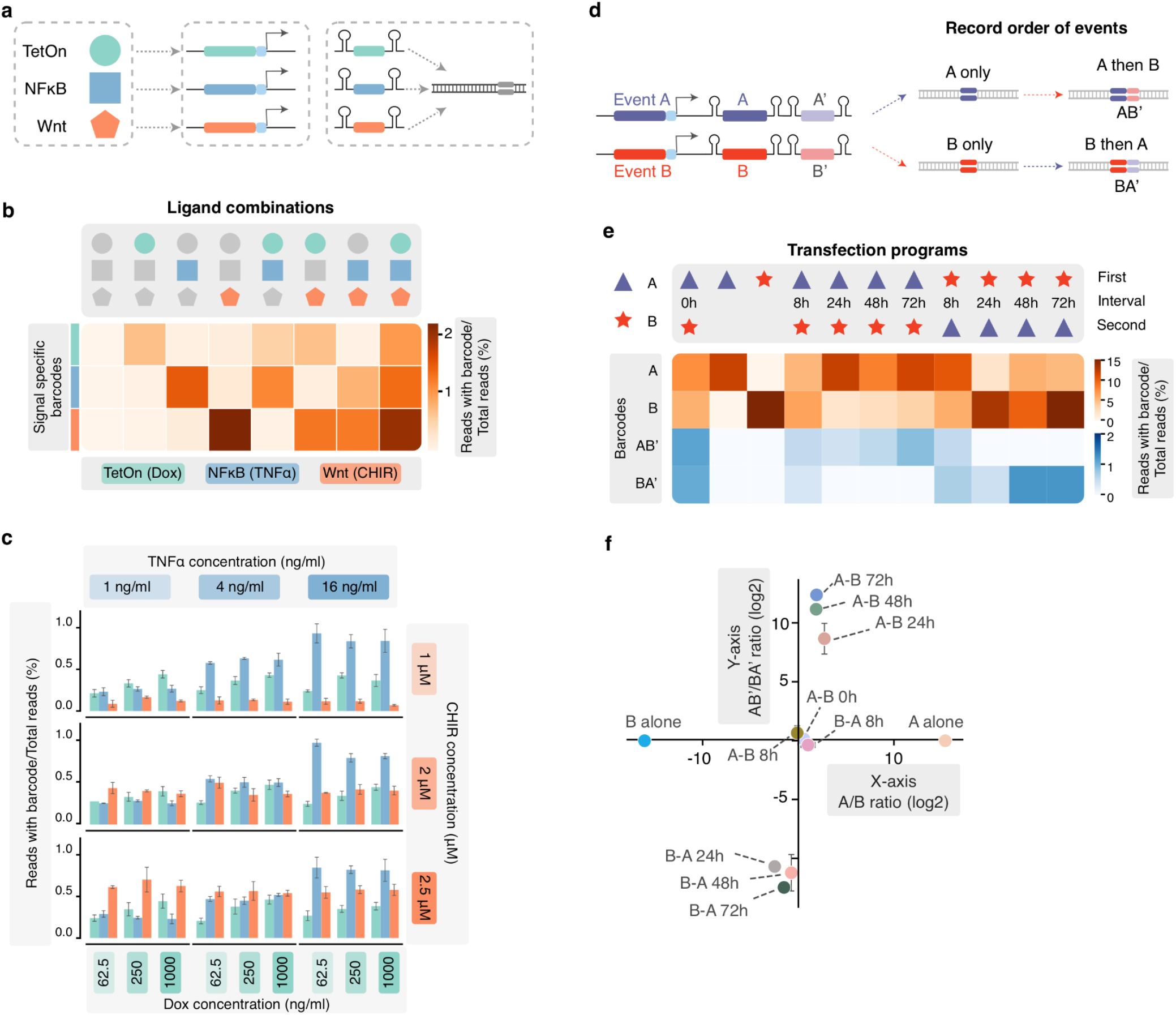
Multiplex recording of signaling pathways or the order of signaling events with ENGRAM. **(a)** Schematic of multiplex recording of signaling pathways. Similar to **Figure 3a** except that all three recorders are integrated within a single population of cells and are writing to a shared DNA Tape. **(b)** Cells bearing multiple recorders were exposed to all possible on/off combinations of three stimuli for 48 hrs, followed by harvesting and sequencing-based quantification of the levels of signal-specific barcodes. Colored shapes as in panel **a**. Concentrations used were 500 ng/ml, 10 ng/ml and 3 µM for doxycycline, TNFα and CHIR99021, respectively. **(c)** Cells bearing multiple recorders were exposed to all possible combinations of high, medium or low concentrations of three stimuli for 48 hrs, followed by harvesting and sequencing-based quantification of the levels of signal-specific barcodes. For Dox, 62.5, 250 or 1000 ng/ml were used; for TNFα, 1, 4 or 16 ng/ml; and for CHIR99021, 1, or 2.5 µM. **(d)** Strategy for ENGRAM-based recording of the order of events A & B. In brief, each signal-responsive recorder programs the expression of two pegRNAs, one of which targets blank DNA Tape, and the other of which targets DNA Tape that has already been edited in response to the other signal. **(e)** We quantified the editing outcomes (A only, B only, A-B’ and B-A’) associated with 11 transfection programs in which either both A & B were introduced simultaneously (1 program), only A or B was introduced (2 programs), or the recorders were serially transfected with varying recovery periods (A→B or B→A; 8 programs). **(f)** The different classes of transfection programs can be distinguished by the ratios of A-B’/B-A’ (y-axis) and A/B editing (x-axis) outcomes. Provided at least 24 hrs of recovery between transfections, A→B programs are readily distinguished from B→A programs. Error bars correspond to standard deviations across 3 transfection replicates.

### Capturing the order in which ENGRAM recorders are active

In the context of a multiplex signal recorder, it is obviously of interest to capture not only the intensity and duration of individual signals, but also the order in which they are active relative to one another. To this end, we devised ENGRAM 2.0 recorders that each comprise an “operon” of multiple, *csy4* hairpin-flanked pegRNAs, each designed to program insertional edits but in a manner that depends on whether other edits had (or had not) already occurred. For example, in the simplest version of this scheme, we might want to map the order of two signaling events, A and B (**Figure 4d**). For this goal, an A-responsive recorder would encode a first pegRNA that that wrote an A-specific barcode to blank DNA Tape (A), but also a second pegRNA that only targeted an already B-edited DNA Tape with a different barcode (A’). Meanwhile, a B-responsive recorder would encode a first pegRNA that that wrote an B-specific barcode to blank DNA Tape (B), but also a second pegRNA that only targeted an already A-edited DNA Tape with a different barcode (B’).

To test this concept, we cloned ENGRAM 2.0 recorders encoding AA’ or BB’ pegRNA operons, each driven by the constitutive PGK promoter. We then performed a series of transfection programs in which either both A & B were introduced simultaneously (1 program), only A or B was introduced (2 programs), or the recorders were serially transfected (A→B or B→A) with the recovery time between transfections varying between 8 and 72 hours (8 programs) (**Figure 4e**). These experiments were performed in triplicate in PE2(+) HEK293T cells, with harvesting, amplification and sequencing of the DNA Tape at five days after the first transfection (11 programs x 3 transfection replicates = 33 conditions) (**Supplementary Figure 4c**). As predicted, and provided there were 24+ hours of recovery between transfections, we observed grossly different ratios of AB’/BA’ edits for (A→B) vs. (B→A) programs (**Figure 4f**; **Supplementary Figure 4d**), indicating that the general scheme is compatible with the recovery of information about the order in which ENGRAM 2.0 recorders are active.

**Supplementary Figure 4.**
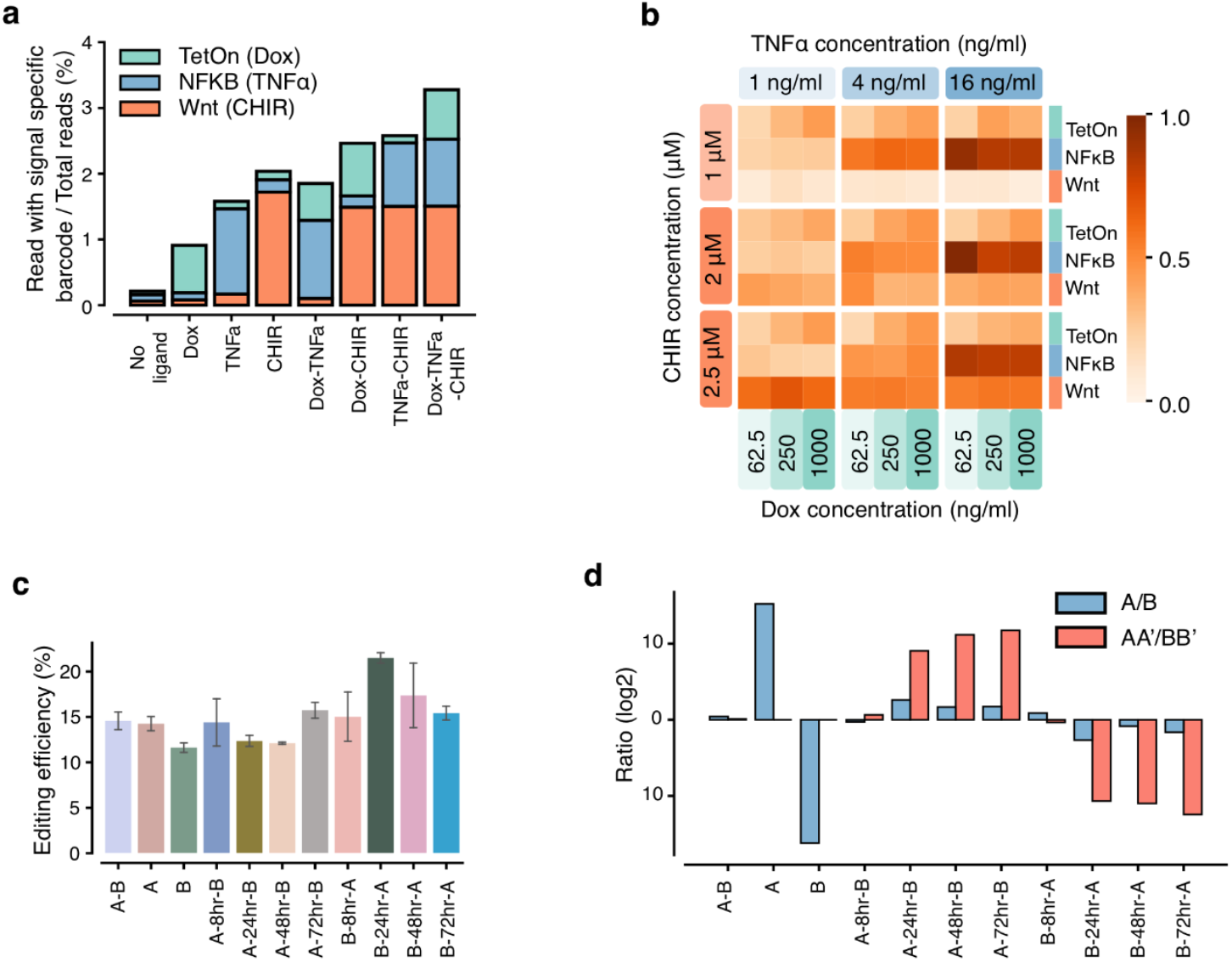
Multiplex recording of signaling pathways or the order of signaling events with ENGRAM. **(a)** Barcode composition of DNA Tape from cells treated with different combinations of stimuli. The recorders exhibit minimal crosstalk between signaling pathways (*e*.*g*. stimulating with CHIR does not lead to appreciable recording by the NF-κB recorder). **(b)** Heatmap visualization of the data shown in **Figure 4c**. Levels of recording are informative of each stimulant’s concentration, even in the context of concurrent recording of three signals to a shared DNA Tape. **(c)** Overall editing efficiencies for the eleven transfection programs represented in **Figure 4e. (d)** Bar plot representation of the same data shown in **Figure 4f**. The different classes of transfection programs can be distinguished by the ratios of A-B’/B-A’ and A/B editing outcomes. Provided at least 24 hrs of recovery between transfections, A→B programs are readily distinguished from B→A programs.

## Discussion

Here we describe ENGRAM, a new strategy for multiplex, DNA-based signal recording, wherein each biological signal of interest is coupled to the Pol-2-mediated transcription of a specific guide RNA, whose expression then programs the insertion of a signal-specific barcode to a genomically encoded DNA Tape. As DNA is stable, recorded signals can be read out at any subsequent point in time, *e*.*g*. by DNA sequencing or, potentially, even by DNA FISH. A key strength of ENGRAM is its multiplexibility. For example, with the 5-bp or 6-bp insertions used here, thousands of distinct biological signals can potentially be recorded within the same cell, all competing to write to a shared DNA Tape. We demonstrate this multiplexibility by showing that analogous to an MPRA, ENGRAM can be applied to concurrently and quantitatively capture the activity of hundreds of enhancers. However, unlike an MPRA, these activities are recorded in the relative abundances of the corresponding insertional barcodes in DNA, rather than being measured from active transcription.

In metazoans, a modest number of core signaling pathways are leveraged to give rise to developmental and functional complexity. To demonstrate how ENGRAM can be applied to record the activity of core signaling pathways, we used Wnt and NF-κB-responsive regulatory elements to drive pegRNAs that write to DNA Tape in a quantitative, specific, signal-responsive manner. We further showed how both the intensity and duration of pathway stimulation contribute to observed levels of recording. We also built and characterized a recorder for Tet-On, highlighting the potential of ENGRAM to be used in conjunction with heterologous signal transduction systems. In a multiplex implementation of these three recorders, there was minimal cross-talk, consistent with the expected orthogonality of these signaling pathways to one another.

Finally, we demonstrated a variant of the ENGRAM method in which the recorder comprises an “operon” of multiple pegRNAs, which are designed to either program or restrict successive edits to the DNA Tape. The resulting pattern of insertional edits allows us to infer the temporal order in which the recorders were activated. Of note, in parallel to this work, we developed a different strategy for “pseudo-processive” genome editing called DNA Ticker Tape. In principle, ENGRAM and DNA Ticker Tape are compatible. For the goal of multiplex, temporally resolved recording of core signaling pathway activity over extended periods of time, the combination of ENGRAM and DNA Ticker Tape may be more powerful than the ENGRAM variant described here.

A number of limitations remain to be addressed. First, as our initial attempts to implement ENGRAM exhibited poor signal-to-noise, we sought to reduce background recording by various means, including by expressing Csy4 as part of the recorder, by introducing auto-regulatory negative feedback on Csy4 levels, by integrating recorder constructs to the genome, and by “blocking” editing with non-targeting pegRNAs during the post-transfection, pre-integration window. Although these strategies were successful, we imagine that further improvements to signal-to-noise might derive from: 1) integrating these and other background reduction strategies; 2) general improvements to prime editing efficiency^24^; and 3) optimization of the specificity of signal-responsive CREs.

Second, here we have only demonstrated ENGRAM recorders for a few hundred enhancer fragments, along with two core (Wnt, NF-κB) and one heterologous (Tet-On) signaling pathway. Looking forward, we envision that many additional such signal-specific recorders can be constructed, validated and optimized. In addition to core signaling pathways, one can also imagine ENGRAM recorders for specific electrical and chemical signals. For pathways for which a signal-responsive, synthetic enhancer is unknown or may not exist, heterologous signal conversion machinery can potentially be constructed and introduced along with the recorder, as we did for Tet-On. Finally, we envision that a set of cell type-specific recorders based on developmental enhancers can be constructed to facilitate recording of the identity of cells’ ancestors.

Third, ENGRAM recorders write to a shared DNA Tape (or DNA Ticker Tape), but each unique recorder is presently ∼1.3 Kb. As such, although transient transfection of a pool of hundreds to thousands of ENGRAM recorders is straightforward, it is more difficult to imagine how dozens or hundreds of recorders can be concurrently integrated to the genome. However, we envision that as additional signal-specific recorders are designed and validated, they can be consolidated to a single “recorder locus”, which can then serve as a common reagent for the multiplex recording of dozens of biological, chemical and electrical signals of interest.

Finally, the deconvolution of ENGRAM signals, especially when recorded to DNA Ticker Tape, will undoubtedly pose some interesting algorithmic challenges. For example, here we show how both the duration and intensity of a signal can contribute to the overall editing rate of a given ENGRAM recorder. If we now imagine recording multiple, fluctuating signals in the context of a dividing and differentiating population of cells, how can we effectively and accurately deconvolve their dynamics?

In summary, ENGRAM is a method for recording specific biological signals to the genome. It is general -- any signal that can be converted to Pol-2 mediated transcription can be used to construct an ENGRAM recorder. It is multiplexable -- by coupling specific signals to specific insertions, the number of signals that can be encoded grows exponentially with the insertion length. It is quantitative -- the strength or duration of signals, and potentially both, can be recorded and recovered. Particularly if combined with DNA Ticker Tape, we envision that ENGRAM can be applied as a means of enriching the DNA-based recordings of cellular histories, across state, space and time.

## Acknowledgements

We thank members of the Shendure Lab for helpful discussions, particularly Xiaoyi Li, Nobuhiko Hamazaki, Jean-Benôit Lalanne, Samuel Regalado, Silvia Domcke, Aaron McKenna, Anna Minkina, Florence Chardon and Troy McDiarmid, as well as members of the Allen Discovery Center for Cell Lineage Tracing, for helpful discussions. We thank David Liu’s lab for sharing the prime editing plasmids. This work was supported by a grant from the Paul G. Allen Frontiers Group (Allen Discovery Center for Cell Lineage Tracing to J.S.) and the National Human Genome Research Institute (UM1HG011586 to J.S.). J.C. is a Howard Hughes Medical Institute Fellow of the Damon Runyon Cancer Research Foundation (DRG-2403-20). J.S. is an Investigator of the Howard Hughes Medical Institute.

## Data availability statement

Raw sequencing data is in the process of being uploaded to the Sequencing Read Archive (SRA) and this preprint will be updated as soon as the link is available. Plasmids for ENGRAM 2.0 recorders are in the process of being deposited to Addgene.

## Materials and Methods

### Cell culture, transient transfections and piggyBAC integrations

HEK293T cells (CRL-11268) and K562 cells (CCL-243) were purchased from ATCC. HEK293T cells and K562 cells were cultured in DMEM High glucose (GIBCO) and RPMI 1640 medium (GIBCO), respectively, supplemented with 10% Fetal Bovine Serum (Rocky Mountain Biologicals) and 1% penicillin-streptomycin (GIBCO). Cells were grown with 5% CO_2_ at 37°C.

For transient transfections, 1 × 10^5^ cells were seeded on a 24-well plate a day before transfection and were transfected with 500 ng plasmid using Lipofectamine 3000 (ThermoFisher L3000015) following the manufacturer’s protocol.

For integrations mediated by the piggyBAC transposon, 1 × 10^5^ cells were seeded on a 24-well plate a day before transfection and then transfected with 500 ng cargo plasmid and 200 ng Super piggyBAC transposase expression vector (SBI) using Lipofectamine 3000 following the manufacturer’s protocol. Monoclonal lines expressing PE2 were constructed by sorting single cells into 96 wells and selected based on prime editing efficiency.

Most ENGRAM recorders tested in this study were integrated into monoclonal PE2(+) HEK293T cell line via the piggyBac transposon method described above. Of note, for doxycycline recorders, an extra integration was performed to introduce reverse tetracycline-controlled transactivator (rtTA), which is activated by doxycycline and binds to the tetracycline response element to activate downstream recorder expression. For recorders co-transfected with blocking pegRNA plasmid, 200 ng plasmid was added to the 500 ng cargo plasmid and 200 ng piggyBac transposase plasmid.

For ligand recording experiments, 1 × 10^5^ cells were seeded on a 48-well plate 6h prior to treatment. 1 ml medium with ligand or negative control was added to each well. For the time series experiment, cells were washed with warm medium and were harvested 24 hours after ligand removal. Doxycycline hyclate (Dox; Sigma) was reconstituted in 1X Phosphate Buffer Solution (PBS) to the final concentration of 10 mg/mL. TNFα (R&D systems, 210-TA-020/CF) was reconstituted in 1 ml PBS to make a 20 µg/ml stock. CHIR-99021 (Selleck, S2924) was purchased as 10 mM stock (1 ml in DMSO). All ligands were stored at - 20°C. Ligands were thawed immediately before experiments, and diluted with the appropriate culturing medium. The same volume of DMSO or PBS was added to the medium as a negative control.

### Library Cloning

The pegRNA-5N recorder (including ENGRAM 1.0, and all three variants of ENGRAM 2.0) was cloned with two steps. First, gene fragment containing CTT pegRNA (Addgene #132778) was PCR amplified using primer sets adding 5-bp degenerate barcode and flanking BsmBI site for the downstream cloning steps. A carrier plasmid containing two BsmBI sites and two *csy4* hairpins was ordered from Twist. Carrier plasmid and the PCR product from the last step were digested with BsmBI (NEB, buffer 3.1) at 55°C for 1h and were purified for ligation. The complete pegRNA with 5N degenerate barcode and *csy4* hairpins was PCR amplified from the ligation product. ENGRAM plasmid and PCR product from above were digested with BsmBI (NEB, buffer 3.1) at 55°C for 1h and purified for ligation. Ligation products were purified and resuspended with 5ul H_2_O for electroporation. Electroporation was performed using NEB® 10-beta Electrocompetent *E. coli* (C3020) with manufacturer’s protocol. Transformed cells were cultured at 30°C overnight.

The libraries of 300 enhancers or plasmids bearing signal-responsive elements were cloned in two steps. First, oligos containing enhancer/CRE, two BsmBI restriction site, barcode, 3’ end of pegRNA and *csy4* hairpin were ordered as oPools from IDT. 5’-ENGRAM 2.0 recorder was digested with Xbal and Ncol (NEB, CutSmart buffer) at 37°C for 1h and purified. Oligos were cloned into the 5’-ENGRAM2.0 recorder using Gibson assembly. Second, a gene fragment containing minP, csy4 hairpin, *HEK3* spacer sequence and pegRNA backbone flanking with two BsmBI sites were ordered as gBlock from IDT. gBlock and construct from step1 were digested with BsmBI (NEB, buffer 3.1) at 55°C for 1h to generate compatible sticky ends and were purified for ligation. Ligation products were transformed into Stable Competent *E*.*coli* (NEB C3040). Transformed cells were cultured at 30°C overnight.

All PCR and digestion purification were purified with AMPure XP beads (0.6x for plasmids and 1.2x for fragments with size 200-300 bp) using manufacturer’s protocol unless specified. All ligation reactions were using Quick ligase (NEB) with vector:insert ratio 1:6 unless specified. All Gibson reactions were using NEBuilder (NEB) with vector:insert ratio 1:6 unless specified. All plasmid DNA was prepared using a ZymoPURE II Plasmid Kit.

### Sequencing Library Generation

Genomic DNA was extracted using protocol as follows: Wash harvested cells with PBS, add 200 µl of freshly prepared lysis buffer (10 mM Tris-HCl, pH 7.5; 0.05% SDS; 25 µg/ml protease (ThermoFisher)) per 0.5-1M cells directly into each well of the tissue culture plate. The genomic DNA mixture was incubated at 50°C for 1 h, followed by an 80°C enzyme inactivation step for 30 min.

For each reaction we used 2 µl of cell lysate, 0.25 µl 100mM forward and reverse primer sets, 22.5 µl H_2_O and 25 µl Robust HotStart ReadyMix 2x (KAPA Biosystems). PCR reactions were performed as follows: 95°C x 3 mins, 22 cycles of (98°C x 20 seconds, 65°C x 15 seconds and 72°C x 40 seconds). The resulting PCR product was then size-selected using a dual size-selection cleanup of 0.5x and 1x AMPure XP beads (Beckman Coulter) to remove genomic DNA and small fragments (< 200 bp) respectively. This size-selected product was subsequently re-amplified to add flow-cell adapter and sample index for 5 cycles. The final PCR product was cleaned with 0.9x AMPure XP beads (Beckman Coulter). The library was sequenced on an Illumina NextSeq 500 sequencer, an Illumina Miseq sequencer, or an Illumina NextSeq 2000 sequencer following manufacturer’s protocol.

### Sequence processing pipeline

Sequences were first aligned to *HEK3* target reference using Burrows-Wheeler Aligner software (*bwa*) with default settings. Aligned reads were then parsed and analyzed for insertion editing efficiencies using pattern-matching functions. For the pool of hexamer barcodes used for enhancer recording, as well as the pentamer barcodes used for signal responsive recording, barcode sequences were chosen to have a Hamming Distance of greater than 2 from all other members of the same set. After extracting barcode sequences from the aligned reads, unexpected barcodes within 1 Hamming Distance from the expected sequences were corrected for insertion counts.

## REFERENCES

1. Li, P. & Elowitz, M. B. Communication codes in developmental signaling pathways. Development 146, (2019).

2. Patwardhan, R. P. et al. High-resolution analysis of DNA regulatory elements by synthetic saturation mutagenesis. Nat. Biotechnol. 27, 1173–1175 (2009).

3. Doupé, D. P. & Perrimon, N. Visualizing and manipulating temporal signaling dynamics with fluorescence-based tools. Sci. Signal. 7, re1 (2014).

4. Inoue, F., Kreimer, A., Ashuach, T., Ahituv, N. & Yosef, N. Identification and Massively Parallel Characterization of Regulatory Elements Driving Neural Induction. Cell Stem Cell 25, 713-727.e10 (2019).

5. Sheth, R. U. & Wang, H. H. DNA-based memory devices for recording cellular events. Nat. Rev. Genet. 19, 718–732 (2018).

6. Tang, W. & Liu, D. R. Rewritable multi-event analog recording in bacterial and mammalian cells. Science 360, (2018).

7. Haurwitz, R. E.,jinek, M., Wiedenheft, B., Zhou, K. & Doudna, J.A. Sequence- and structure-specific RNA processing by a CRISPR endonuclease. Science 329, 1355–1358 (2010).

8. Sternberg, S. H., Haurwitz, R. E. & Doudna, J. A. Mechanism of substrate selection by a highly specific CRISPR endoribonuclease. RNA 18, 661–67 (2012).

9. Haurwitz, R. E., Sternberg, S. H. & Doudna, J. A. Csy4 relies on an unusual catalytic dyad to position and cleave CRISPR RNA. The EMBO Journal vol. 31 2824–2832 (2012).

10. Nissim, L., Perli, S. D., Fridkin, A., Perez-Pinera, P. & Lu, T. K. Multiplexed and programmable regulation of gene networks with an integrated RNA and CRISPR/Cas toolkit in human cells. Mol. Cell 54, 698–710 (2014).

11. Anzalone, A. V. et al. Search-and-replace genome editing without double-strand breaks or donor DNA. Nature 576, 149–157 (2019).

12. Shipman, S. L., Nivala, J., Macklis, J. D. & Church, G. M. Molecular recordings by directed CRISPR spacer acquisition. Science 353, aaf1175 (2016).

13. Shipman, S. L., Nivala, J., Macklis, J. D. & Church, G. M. CRISPR-Cas encoding of a digital movie into the genomes of a population of living bacteria. Nature 547, 345–349 (2017).

14. Sheth, R. U., Yim, S. S., Wu, F. L. & Wang, H. H. Multiplex recording of cellular events over time on CRISPR biological tape. Science 358, 1457–1461 (2017).

15. Schmidt, F., Cherepkova, M. Y. & Platt, R. J. Transcriptional recording by CRISPR spacer acquisition from RNA. Nature 562, 380–385 (2018).

16. Yim, S. S. et al. Robust direct digital-to-biological data storage in living cells. Nat. Chem. Biol. 17, 246–253 (2021).

17. Bhattarai-Kline, S. et al. Reconstructing transcriptional histories by CRISPR acquisition of retron-based genetic barcodes. bioRxiv 2021.08.11.455990 (2021) doi:10.1101/2021.08.11.455990.

18. Klein, J. C. et al. A systematic evaluation of the design and context dependencies of massively parallel reporter assays. Nat. Methods 17, 1083–1091 (2020).

19. Raj, B. et al. Simultaneous single-cell profiling of lineages and cell types in the vertebrate brain. Nat. Biotechnol. 36, 442–450 (2018).

20. Muerdter, F. et al. Resolving systematic errors in widely used enhancer activity assays in human cells. Nat. Methods 15, 141–149 (2018).

21. Gossen, M. et al. Transcriptional Activation by Tetracyclines in Mammalian Cells. Science vol. 268 1766–1769 (1995).

22. Zabel, U., Schreck, R. & Baeuerle, P. A. DNA binding of purified transcription factor NF-kappa B. Affinity, specificity, Zn2 dependence, and differential half-site recognition. Journal of Biological Chemistry vol. 266 252 – 260 (1991).

23. pGL4.49[luc2P/TCF-LEF/Hygro] Vector Protocol. https://www.promega.com/resources/protocols/product-information-sheets/a/pgl4-49-vector-protocol/.

24. Chen, P. . et al. Enhanced prime editing systems by manipulating cellular determinants of editing outcomes. Cell 184, 5635-5652.e29 (2021).

